## Supplementary figures and images for "The protein expression profile of ACE2 in human tissues"

### Supplementary Figure 1

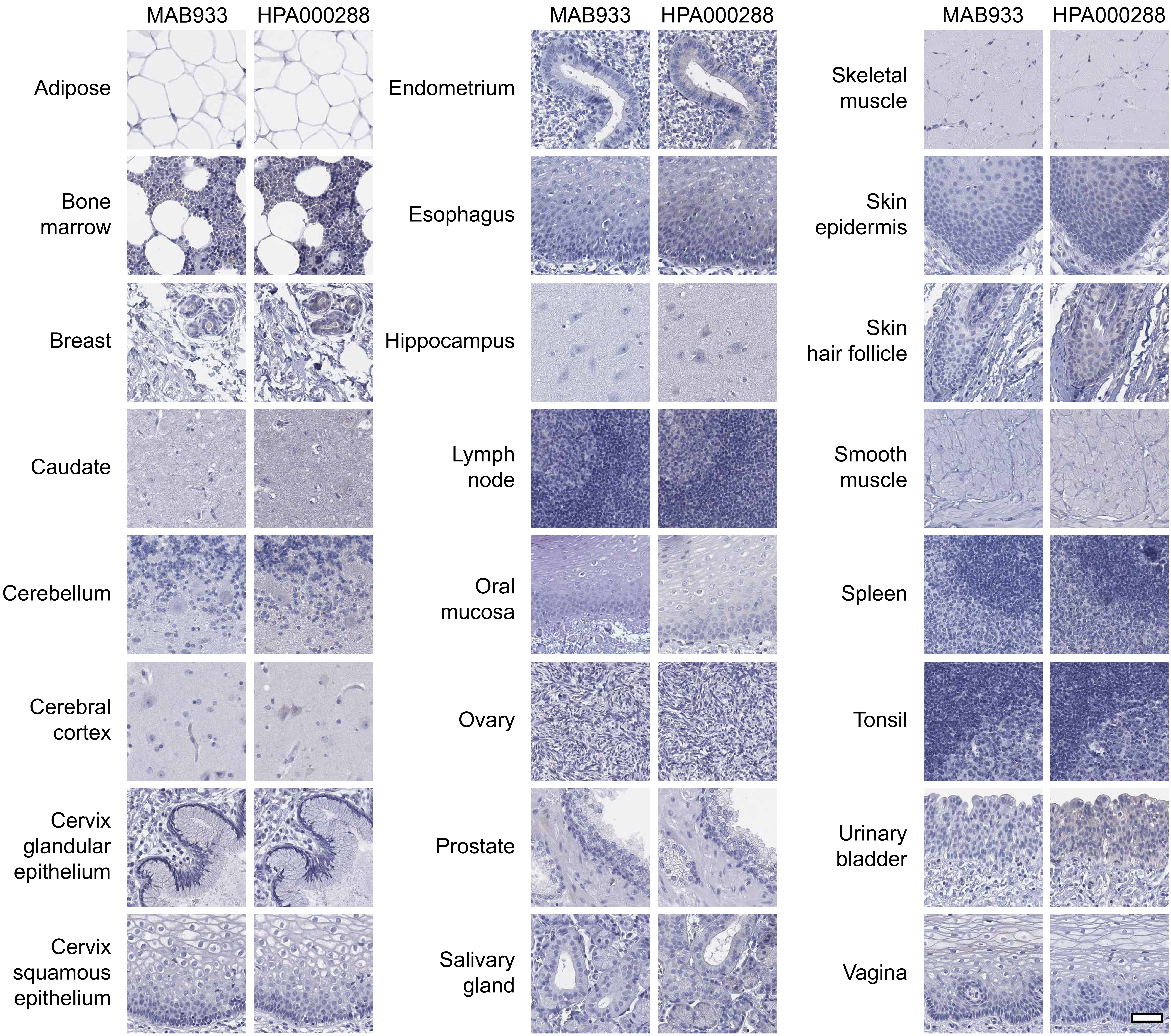
