## Supplementary Table 1 for "The protein expression profile of ACE2 in human tissues"

**Table Expanded View 1. Number of samples used for immunohistochemical analysis of each normal tissue type, including information on age and gender.**

| MAB933<br>Tissue | Total samples per tissue | Large sections | TMA samples | Female | Male | 0-9 years | 10-19 years | 20-29 years | 30-39 years | 40-49 years | 50-59 years | 60-69 years | 70-79 years | 80+ years |
|---|---|---|---|---|---|---|---|---|---|---|---|---|---|---|
| adipose tissue | 10 | 0 | 10 | 2 | 8 | 0 | 1 | 2 | 1 | 0 | 2 | 2 | 2 | 0 |
| adrenal gland | 5 | 0 | 5 | 2 | 3 | 0 | 0 | 0 | 0 | 1 | 3 | 1 | 0 | 0 |
| appendix | 8 | 0 | 8 | 4 | 4 | 1 | 1 | 2 | 0 | 2 | 1 | 0 | 1 | 0 |
| bone marrow | 9 | 0 | 9 | 3 | 6 | 0 | 0 | 0 | 0 | 1 | 2 | 4 | 2 | 0 |
| breast | 5 | 0 | 5 | 5 | 0 | 0 | 0 | 1 | 2 | 2 | 0 | 0 | 0 | 0 |
| bronchioli | 6 | 6 | 0 | 4 | 2 | 0 | 0 | 0 | 0 | 1 | 1 | 2 | 2 | 0 |
| bronchus | 13 | 8 | 5 | 6 | 7 | 0 | 0 | 0 | 0 | 1 | 2 | 7 | 3 | 0 |
| caudate | 8 | 0 | 8 | 3 | 5 | 0 | 1 | 0 | 1 | 1 | 2 | 1 | 2 | 0 |
| cerebellum | 8 | 0 | 8 | 5 | 3 | 0 | 2 | 1 | 1 | 0 | 2 | 1 | 1 | 0 |
| cerebral cortex | 7 | 0 | 7 | 1 | 6 | 0 | 0 | 0 | 1 | 0 | 3 | 1 | 2 | 0 |
| cervix, uterine | 6 | 0 | 6 | 6 | 0 | 0 | 0 | 1 | 3 | 1 | 1 | 0 | 0 | 0 |
| colon | 7 | 0 | 7 | 4 | 3 | 0 | 1 | 0 | 0 | 0 | 0 | 3 | 1 | 2 |
| duodenum | 6 | 0 | 6 | 3 | 3 | 0 | 0 | 0 | 0 | 0 | 3 | 2 | 1 | 0 |
| endometrium | 17 | 0 | 17 | 17 | 0 | 0 | 0 | 1 | 7 | 6 | 2 | 1 | 0 | 0 |
| epididymis | 10 | 0 | 10 | 0 | 10 | 0 | 0 | 3 | 2 | 2 | 2 | 0 | 1 | 0 |
| esophagus | 6 | 0 | 6 | 2 | 4 | 0 | 0 | 0 | 0 | 0 | 2 | 1 | 1 | 2 |
| eye | 1 | 1 | 0 | 0 | 1 | 0 | 0 | 0 | 0 | 0 | 0 | 1 | 0 | 0 |
| fallopian tube | 11 | 0 | 11 | 11 | 0 | 0 | 0 | 2 | 5 | 2 | 2 | 0 | 0 | 0 |
| gallbladder | 8 | 0 | 8 | 4 | 4 | 0 | 0 | 1 | 2 | 1 | 1 | 3 | 0 | 0 |
| heart muscle | 9 | 0 | 9 | 3 | 6 | 0 | 0 | 0 | 1 | 1 | 5 | 2 | 0 | 0 |
| hippocampus | 8 | 0 | 8 | 2 | 6 | 0 | 1 | 1 | 0 | 2 | 3 | 0 | 1 | 0 |
| kidney | 7 | 0 | 7 | 2 | 5 | 0 | 1 | 0 | 0 | 1 | 2 | 1 | 2 | 0 |
| liver | 8 | 0 | 8 | 5 | 3 | 0 | 0 | 0 | 1 | 0 | 4 | 2 | 1 | 0 |
| lung | 367 | 0 | 367 | 186 | 181 | 0 | 0 | 1 | 0 | 11 | 45 | 168 | 130 | 12 |
| lymph node | 11 | 0 | 11 | 7 | 4 | 0 | 0 | 1 | 2 | 0 | 7 | 1 | 0 | 0 |
| nasopharynx | 17 | 12 | 5 | 2 | 6 | 0 | 2 | 2 | 1 | 1 | 5 | 3 | 2 | 1 |
| oral mucosa | 10 | 0 | 10 | 6 | 4 | 0 | 0 | 0 | 0 | 0 | 2 | 4 | 1 | 3 |
| ovary | 7 | 0 | 7 | 7 | 0 | 0 | 0 | 1 | 4 | 2 | 0 | 0 | 0 | 0 |
| pancreas | 10 | 0 | 10 | 6 | 4 | 0 | 0 | 0 | 1 | 3 | 1 | 3 | 2 | 0 |
| parathyroid gland | 4 | 0 | 4 | 1 | 3 | 0 | 0 | 0 | 0 | 1 | 1 | 0 | 0 | 2 |
| placenta | 7 | 0 | 7 | 7 | 0 | 0 | 2 | 1 | 4 | 0 | 0 | 0 | 0 | 0 |
| prostate | 9 | 0 | 9 | 0 | 9 | 0 | 0 | 0 | 1 | 0 | 1 | 5 | 2 | 0 |
| rectum | 9 | 0 | 9 | 4 | 5 | 0 | 0 | 0 | 0 | 1 | 3 | 4 | 1 | 0 |
| salivary gland | 7 | 0 | 7 | 3 | 4 | 0 | 0 | 1 | 2 | 1 | 0 | 2 | 1 | 0 |
| seminal vesicle | 8 | 0 | 8 | 0 | 8 | 0 | 0 | 0 | 1 | 1 | 3 | 2 | 1 | 0 |
| skeletal muscle | 6 | 0 | 6 | 2 | 4 | 0 | 0 | 0 | 1 | 0 | 2 | 1 | 1 | 1 |
| skin | 15 | 0 | 15 | 7 | 8 | 0 | 1 | 2 | 1 | 1 | 3 | 2 | 5 | 0 |

| | | | | | | | | | | | | | | |
|---|---|---|---|---|---|---|---|---|---|---|---|---|---|---|
| small intestine | 8 | 0 | 8 | 5 | 3 | 1 | 0 | 0 | 1 | 1 | 2 | 1 | 0 | 2 |
| smooth muscle | 9 | 0 | 9 | 5 | 4 | 0 | 2 | 0 | 2 | 1 | 1 | 1 | 0 | 2 |
| spleen | 7 | 0 | 7 | 2 | 5 | 0 | 2 | 0 | 0 | 0 | 2 | 1 | 2 | 0 |
| stomach | 15 | 0 | 15 | 5 | 10 | 0 | 0 | 0 | 0 | 2 | 6 | 3 | 4 | 0 |
| testis | 8 | 0 | 8 | 0 | 8 | 0 | 0 | 3 | 1 | 2 | 1 | 1 | 0 | 0 |
| thyroid gland | 8 | 0 | 8 | 5 | 3 | 0 | 0 | 2 | 1 | 1 | 2 | 2 | 0 | 0 |
| tonsil | 10 | 0 | 10 | 6 | 4 | 1 | 5 | 0 | 3 | 1 | 0 | 0 | 0 | 0 |
| urinary bladder | 7 | 0 | 7 | 2 | 5 | 0 | 0 | 0 | 0 | 1 | 0 | 3 | 2 | 1 |
| vagina | 8 | 0 | 8 | 8 | 0 | 0 | 0 | 0 | 2 | 5 | 0 | 1 | 0 | 0 |
| **GRAND TOTAL** | 750 | 27 | 723 | 370 | 371 | 3 | 22 | 29 | 55 | 61 | 132 | 243 | 177 | 28 |

**HPA000288**

| Tissue | Total samples per tissue | Large sections | TMA samples | Female | Male | 0-9 years | 10-19 years | 20-29 years | 30-39 years | 40-49 years | 50-59 years | 60-69 years | 70-79 years | 80+ years |
|---|---|---|---|---|---|---|---|---|---|---|---|---|---|---|
| adipose tissue | 13 | 0 | 13 | 3 | 10 | 0 | 1 | 1 | 1 | 2 | 4 | 3 | 1 | 0 |
| adrenal gland | 5 | 0 | 5 | 2 | 3 | 0 | 0 | 0 | 0 | 1 | 3 | 1 | 0 | 0 |
| appendix | 10 | 0 | 10 | 6 | 4 | 0 | 2 | 3 | 1 | 2 | 1 | 0 | 1 | 0 |
| bone marrow | 11 | 0 | 11 | 4 | 7 | 0 | 0 | 0 | 0 | 1 | 2 | 3 | 4 | 1 |
| breast | 7 | 0 | 7 | 7 | 0 | 0 | 0 | 0 | 2 | 4 | 1 | 0 | 0 | 0 |
| bronchioli | 6 | 6 | 0 | 4 | 2 | 0 | 0 | 0 | 0 | 1 | 1 | 2 | 2 | 0 |
| bronchus | 13 | 8 | 5 | 6 | 7 | 0 | 0 | 0 | 0 | 1 | 2 | 7 | 3 | 0 |
| caudate | 10 | 0 | 10 | 4 | 6 | 0 | 1 | 0 | 1 | 2 | 3 | 1 | 2 | 0 |
| cerebellum | 8 | 0 | 8 | 5 | 3 | 0 | 2 | 1 | 0 | 1 | 2 | 1 | 1 | 0 |
| cerebral cortex | 10 | 0 | 10 | 3 | 7 | 0 | 1 | 0 | 1 | 1 | 4 | 1 | 2 | 0 |
| cervix, uterine | 8 | 0 | 8 | 8 | 0 | 0 | 0 | 1 | 2 | 1 | 2 | 0 | 1 | 1 |
| colon | 8 | 0 | 8 | 4 | 4 | 1 | 0 | 0 | 0 | 0 | 2 | 0 | 2 | 2 |
| duodenum | 8 | 0 | 8 | 5 | 3 | 0 | 0 | 0 | 0 | 1 | 2 | 2 | 3 | 0 |
| endometrium | 22 | 0 | 22 | 22 | 0 | 0 | 0 | 1 | 10 | 6 | 4 | 0 | 0 | 1 |
| epididymis | 7 | 0 | 7 | 0 | 7 | 0 | 0 | 2 | 2 | 1 | 2 | 0 | 0 | 0 |
| esophagus | 9 | 0 | 9 | 3 | 6 | 0 | 0 | 0 | 0 | 0 | 2 | 3 | 2 | 2 |
| eye | 1 | 1 | 0 | 0 | 1 | 0 | 0 | 0 | 0 | 0 | 0 | 1 | 0 | 0 |
| fallopian tube | 12 | 0 | 12 | 12 | 0 | 0 | 0 | 2 | 5 | 3 | 2 | 0 | 0 | 0 |
| gallbladder | 9 | 0 | 9 | 5 | 4 | 0 | 0 | 1 | 0 | 2 | 1 | 3 | 2 | 0 |
| heart muscle | 9 | 0 | 9 | 4 | 5 | 0 | 1 | 0 | 1 | 0 | 5 | 2 | 0 | 0 |
| hippocampus | 9 | 0 | 9 | 2 | 7 | 0 | 1 | 1 | 0 | 3 | 3 | 0 | 1 | 0 |
| kidney | 9 | 0 | 9 | 3 | 6 | 0 | 1 | 0 | 1 | 1 | 3 | 2 | 1 | 0 |
| liver | 10 | 0 | 10 | 6 | 4 | 0 | 0 | 0 | 3 | 0 | 4 | 1 | 2 | 0 |
| lung | 370 | 0 | 370 | 188 | 182 | 0 | 0 | 1 | 0 | 11 | 45 | 169 | 132 | 12 |
| lymph node | 12 | 0 | 12 | 7 | 5 | 0 | 0 | 0 | 3 | 0 | 7 | 2 | 0 | 0 |
| nasopharynx | 17 | 12 | 5 | 3 | 14 | 1 | 1 | 1 | 1 | 0 | 5 | 4 | 3 | 1 |
| oral mucosa | 8 | 0 | 8 | 5 | 3 | 0 | 0 | 0 | 0 | 0 | 2 | 3 | 1 | 2 |
| ovary | 8 | 0 | 8 | 8 | 0 | 0 | 0 | 1 | 3 | 2 | 0 | 1 | 1 | 0 |

| | | | | | | | | | | | | | | |
|---|---|---|---|---|---|---|---|---|---|---|---|---|---|---|
| pancreas | 11 | 0 | 11 | 6 | 5 | 0 | 0 | 0 | 1 | 2 | 2 | 2 | 4 | 0 |
| parathyroid gland | 4 | 0 | 4 | 2 | 2 | 0 | 0 | 0 | 0 | 0 | 1 | 0 | 1 | 2 |
| placenta | 9 | 0 | 9 | 9 | 0 | 0 | 2 | 2 | 4 | 1 | 0 | 0 | 0 | 0 |
| prostate | 11 | 0 | 11 | 0 | 11 | 0 | 0 | 0 | 1 | 0 | 1 | 7 | 2 | 0 |
| rectum | 9 | 0 | 9 | 4 | 5 | 0 | 0 | 0 | 0 | 1 | 2 | 4 | 2 | 0 |
| salivary gland | 10 | 0 | 10 | 4 | 6 | 0 | 0 | 1 | 2 | 1 | 2 | 3 | 1 | 0 |
| seminal vesicle | 9 | 0 | 9 | 0 | 9 | 0 | 0 | 0 | 1 | 1 | 1 | 5 | 1 | 0 |
| skeletal muscle | 7 | 0 | 7 | 2 | 5 | 0 | 0 | 0 | 1 | 0 | 2 | 1 | 2 | 1 |
| skin | 15 | 0 | 15 | 6 | 9 | 0 | 0 | 0 | 1 | 2 | 3 | 2 | 4 | 3 |
| small intestine | 8 | 0 | 8 | 5 | 3 | 1 | 0 | 0 | 2 | 1 | 2 | 1 | 0 | 1 |
| smooth muscle | 9 | 0 | 9 | 5 | 4 | 0 | 1 | 0 | 2 | 2 | 1 | 2 | 0 | 1 |
| spleen | 8 | 0 | 8 | 2 | 6 | 0 | 3 | 0 | 0 | 0 | 2 | 1 | 2 | 0 |
| stomach | 16 | 0 | 16 | 6 | 10 | 0 | 0 | 0 | 0 | 2 | 5 | 3 | 5 | 1 |
| testis | 10 | 0 | 10 | 0 | 10 | 0 | 0 | 4 | 2 | 2 | 1 | 1 | 0 | 0 |
| thyroid gland | 9 | 0 | 9 | 4 | 5 | 0 | 0 | 1 | 2 | 0 | 3 | 2 | 1 | 0 |
| tonsil | 9 | 0 | 9 | 6 | 3 | 2 | 4 | 0 | 2 | 1 | 0 | 0 | 0 | 0 |
| urinary bladder | 6 | 0 | 6 | 2 | 4 | 0 | 0 | 0 | 0 | 0 | 0 | 2 | 3 | 1 |
| vagina | 9 | 0 | 9 | 9 | 0 | 0 | 0 | 0 | 1 | 6 | 0 | 2 | 0 | 0 |
| **GRAND TOTAL** | 798 | 27 | 771 | 401 | 397 | 5 | 21 | 24 | 59 | 69 | 142 | 250 | 195 | 32 |
