## Supplementary Table 2 for "The protein expression profile of ACE2 in human tissues"

**Table Expanded View 2.** Summary of protein expression patterns in 156 different cell types and 45 tissues based on two antibodies, graded as not detected (0), low (1), medium (2) or high (3), taking into consideration immunohistochemical staining intensity and quantity of positive cells.

| Cell type | MAB933 | HPA000288 |
|---|---|---|
| Small intestine - enterocytes | 3 | 3 |
| Small intestine - goblet cells | 0 | 0 |
| Small intestine - endocrine cells | 0 | 0 |
| Small intestine - Paneth cells | 0 | 0 |
| Small intestine - crypt cells | 2 | 2 |
| Small intestine - endothelial cells/pericytes | 0 | 0 |
| Colon - enterocytes | 2 | 2 |
| Colon - goblet cells | 0 | 0 |
| Colon - endocrine cells | 0 | 0 |
| Colon - crypt cells | 0 | 0 |
| Colon - endothelial cells/pericytes | 0 | 0 |
| Colon - peripheral nerve | 0 | 0 |
| Duodenum - enterocytes | 3 | 3 |
| Duodenum - goblet cells | 0 | 0 |
| Duodenum - endocrine cells | 0 | 0 |
| Duodenum - crypt cells | 2 | 2 |
| Duodenum - glands of Brunner | 1 | 1 |
| Duodenum - endothelial cells/pericytes | 0 | 0 |
| Kidney - proximal tubule cells | 3 | 3 |
| Kidney - distal tubule cells | 0 | 0 |
| Kidney - collecting duct cells | 0 | 0 |
| Kidney - Bowman's capsule | 1 | 2 |
| Kidney - podocytes | 0 | 0 |
| Kidney - mesangial cells | 0 | 0 |
| Kidney - endothelial cells/pericytes | 0 | 0 |
| Testis - Sertoli cells | 3 | 3 |
| Testis - Leydig cells | 3 | 3 |
| Testis - endothelial cells/pericytes | 0 | 0 |
| Testis - peritubular cells | 0 | 0 |
| Testis - spermatogonia | 0 | 0 |
| Testis - prelep. spermatocytes | 0 | 0 |
| Testis - pachy. spermatocytes | 0 | 0 |
| Testis - early spermatids | 0 | 0 |
| Testis - late spermatids | 0 | 0 |
| Gallbladder - endothelial cells/pericytes | 0 | 0 |
| Gallbladder - glandular cells | 3 | 3 |
| Heart - cardiomyocytes | 1 | 2 |
| Heart muscle - endothelial cells/pericytes | 1 | 1 |
| Thyroid gland - glandular cells | 0 | 0 |
| Thyroid gland - endothelial cells/pericytes | 1 | 1 |
| Adipose tissue - adipocytes | 0 | 0 |
| Epididymis - glandular cells | 1 | 1 |
| Epididymis - endothelial cells/pericytes | 0 | 0 |
| Breast - glandular cells | 0 | 0 |
| Breast - myoepithelial cells | 0 | 0 |
| Breast - adipocytes | 0 | 0 |

| | | |
|---|---|---|
| Pancreas - islets of Langerhans | 0 | 0 |
| Pancreas - acinar glandular cells | 0 | 0 |
| Pancreas - intercalated ducts | 0 | 0 |
| Pancreas - intralobular ducts | 0 | 0 |
| Pancreas - interlobular ducts | 2 | 2 |
| Pancreas - endothelial cells/pericytes | 1 | 1 |
| Rectum - enterocytes | 1 | 1 |
| Rectum - goblet cells | 0 | 0 |
| Rectum - endocrine cells | 0 | 0 |
| Rectum - endothelial cells/pericytes | 0 | 0 |
| Ovary - follicle cells | 0 | 0 |
| Ovary - stromal cells | 0 | 0 |
| Esophagus - squamous epithelial cells | 0 | 0 |
| Oral mucosa - squamous epithelial cells | 0 | 0 |
| Liver - hepatocytes | 0 | 0 |
| Liver - bile duct cells | 1 | 1 |
| Liver - Kupffer cells | 0 | 0 |
| Liver - endothelial cells/pericytes | 0 | 0 |
| Seminal vesicle - glandular cells | 2 | 2 |
| Seminal vesicle - endothelial cells/pericytes | 0 | 0 |
| Salivary gland - serous acinar cells | 0 | 0 |
| Salivary gland - mucinous acinar cells | 0 | 0 |
| Salivary gland - ductal cells | 0 | 0 |
| Salivary gland - endothelial cells/pericytes | 0 | 0 |
| Placenta - syncytiotrophoblasts | 1 | 2 |
| Placenta - cytotrophoblasts | 1 | 2 |
| Placenta - extravillous trophoblasts | 1 | 2 |
| Placenta - decidual cells | 0 | 0 |
| Placenta - endothelial cells/pericytes | 0 | 0 |
| Vagina - squamous epithelial cells | 0 | 0 |
| Nasal mucosa - ciliated cells | 0 | 1 |
| Nasal mucosa - goblet cells | 0 | 0 |
| Nasal mucosa - basal cells | 0 | 0 |
| Nasal mucosa - squamous epithelial cells | 0 | 0 |
| Nasal mucosa - submucosal glands | 0 | 0 |
| Nasal mucosa - endothelial cells/pericytes | 0 | 0 |
| Bronchus - ciliated cells | 0 | 1 |
| Bronchus - goblet cells | 0 | 0 |
| Bronchus - basal cells | 0 | 0 |
| Bronchus - endothelial cells/pericytes | 0 | 0 |
| Bronchus - submucosal glands | 0 | 0 |
| Lung - alveolar cells type I | 0 | 0 |
| Lung - alveolar cells type II | 0 | 0 |
| Lung - bronchioli | 0 | 0 |
| Lung - macrophages | 0 | 0 |
| Lung - endothelial cells/pericytes | 0 | 0 |
| Appendix - enterocytes | 1 | 1 |
| Appendix - goblet cells | 0 | 0 |
| Appendix - endocrine cells | 0 | 0 |
| Appendix - lymphoid tissue | 0 | 0 |
| Appendix - endothelial cells/pericytes | 0 | 0 |
| Skeletal muscle - myocytes | 0 | 0 |
| Fallopian tube - ciliated cells | 0 | 1 |

| | | |
|---|---|---|
| Fallopian tube - non-ciliated cells | 0 | 0 |
| Fallopian tube - endothelial cells/pericytes | 1 | 1 |
| Lymph node - germinal center cells | 0 | 0 |
| Lymph node - non-germinal center cells | 0 | 0 |
| Stomach corpus - surface epithelial cells | 0 | 0 |
| Stomach corpus - parietal cells | 0 | 0 |
| Stomach corpus - chief cells | 0 | 0 |
| Stomach corpus - endothelial cells/pericytes | 0 | 0 |
| Stomach antrum - surface epithelial cells | 1 | 1 |
| Stomach antrum - endothelial cells/pericytes | 0 | 0 |
| Prostate - glandular cells | 0 | 0 |
| Prostate - endothelial cells/pericytes | 0 | 0 |
| Adrenal gland - glandular cells | 0 | 0 |
| Adrenal gland - endothelial cells/pericytes | 1 | 1 |
| Endometrium - glandular cells | 0 | 0 |
| Endometrium - stromal cells | 0 | 0 |
| Urinary bladder - urothelial cells | 0 | 0 |
| Cervix, uterine - glandular cells | 0 | 0 |
| Cervix, uterine - squamous epithelial cells | 0 | 0 |
| Smooth muscle - smooth muscle cells | 0 | 0 |
| Cerebral cortex - neuronal cells | 0 | 0 |
| Cerebral cortex - glial cells | 0 | 0 |
| Cerebral cortex - neuropil | 0 | 0 |
| Cerebral cortex - endothelial cells/pericytes | 0 | 0 |
| Cerebellum - cells in granular layer | 0 | 0 |
| Cerebellum - cells in molecular layer | 0 | 0 |
| Cerebellum - Purkinje cells | 0 | 0 |
| Hippocampus - neuronal cells | 0 | 0 |
| Hippocampus - glial cells | 0 | 0 |
| Caudate - neuronal cells | 0 | 0 |
| Caudate - glial cells | 0 | 0 |
| Eye - cornea | 1 | 1 |
| Eye - conjunctiva | 2 | 2 |
| Eye - hyaloid membrane | 0 | 0 |
| Eye - lens epithelial cells | 0 | 0 |
| Eye - lens fiber cells | 0 | 0 |
| Eye - inner nuclear layer | 0 | 0 |
| Eye - rod photoreceptor cells | 0 | 0 |
| Eye - cone photoreceptor cells | 0 | 0 |
| Eye - ganglion cells | 0 | 0 |
| Eye - limiting membrane | 0 | 0 |
| Eye - inner plexiform layer | 0 | 0 |
| Eye - nerve fiber layer | 0 | 0 |
| Eye - outer plexiform layer | 0 | 0 |
| Eye - pigment epithelium | 0 | 0 |
| Eye - endothelial cells/pericytes | 0 | 0 |
| Skin - keratinocytes | 0 | 0 |
| Skin - melanocytes | 0 | 0 |
| Skin - Langerhans | 0 | 0 |
| Skin - hair follicle | 0 | 0 |
| Skin - eccrine glands | 0 | 0 |
| Skin - fibroblasts | 0 | 0 |
| Spleen - cells in white pulp | 0 | 0 |

| | | |
|---|---|---|
| Spleen - cells in red pulp | 0 | 0 |
| Tonsil - germinal center cells | 0 | 0 |
| Tonsil - non-germinal center cells | 0 | 0 |
| Tonsil - squamous epithelial cells | 0 | 0 |
| Parathyroid gland - glandular cells | 0 | 0 |
| Parathyroid gland - endothelial cells/pericytes | 1 | 1 |
| Bone marrow - hematopoietic cells | 0 | 0 |
