## Supplementary Table 3 for "The protein expression profile of ACE2 in human tissues"

**Table Expanded View 3. List of samples included in the cohort of normal lung tissues (n=360), with information on age, gender, smoking status ,and performance status, and immunohistochemical staining of ACE2.**

* = 0, Fully active, able to carry on all pre-disease performance without restriction; 1, Restricted in physically strenuous activity but ambulatory and able to carry out work of a light or sedentary nature, e.g., light house work, office work; 2, Ambulatory and capable of all selfcare but unable to carry out any work activities. Up and about more than 50% of waking hours; 3, Capable of only limited selfcare, confined to bed or chair more than 50% of waking hours; 4, Completely disabled. Cannot carry on any selfcare. Totally confined to bed or chair.

| Individual | Age (years) | Gender (F=female, M=male) | Smoking (C=current, F=former, N=never) | Performance status (ECOG/WHO)* | Positive (+) or negative (-) IHC staining MAB933 | Positive (+) or negative (-) IHC staining HPA000288 |
|---|---|---|---|---|---|---|
| 1 | 78 | M | N | 1 | - | - |
| 2 | 74 | M | F | 1 | - | - |
| 3 | 74 | F | C | 0 | - | - |
| 4 | 73 | M | C | 1 | - | - |
| 5 | 74 | F | F | 0 | - | - |
| 6 | 64 | M | N | 0 | - | - |
| 7 | 64 | M | C | 0 | - | - |
| 8 | 63 | F | F | 0 | - | - |
| 9 | 61 | F | C | 0 | - | - |
| 10 | 58 | F | N | 0 | - | - |
| 11 | 83 | F | F | 1 | - | - |
| 12 | 72 | F | F | 0 | - | - |
| 13 | 71 | M | C | 0 | - | - |
| 14 | 70 | F | N | 0 | - | - |
| 15 | 70 | M | F | 0 | - | - |
| 16 | 65 | M | C | 0 | - | - |
| 17 | 61 | F | F | 0 | - | - |
| 18 | 57 | F | C | 1 | - | - |
| 19 | 47 | M | C | 1 | - | - |
| 20 | 70 | M | F | 1 | - | - |
| 21 | 80 | M | F | 1 | - | - |
| 22 | 80 | M | F | 0 | - | - |
| 23 | 78 | M | C | 1 | - | - |
| 24 | 75 | F | C | 1 | - | - |
| 25 | 74 | F | F | 1 | - | - |
| 26 | 74 | F | C | 2 | - | - |
| 27 | 71 | F | F | 0 | - | - |
| 28 | 67 | M | F | 0 | - | - |
| 29 | 65 | M | C | 1 | - | - |
| 30 | 65 | M | C | 0 | - | - |
| 31 | 64 | M | C | 1 | - | - |
| 32 | 63 | M | C | 2 | - | - |
| 33 | 62 | M | F | 0 | - | - |
| 34 | 58 | F | N | 0 | - | - |
| 35 | 52 | F | C | 0 | - | - |
| 36 | 48 | M | C | 1 | - | - |
| 37 | 46 | F | C | 0 | - | - |
| 38 | 76 | F | C | 0 | - | - |
| 39 | 75 | M | N | 0 | - | - |
| 40 | 70 | M | C | 0 | - | - |
| 41 | 67 | M | F | 0 | - | - |
| 42 | 67 | F | F | 0 | - | - |
| 43 | 66 | M | F | 0 | - | - |
| 44 | 65 | F | C | 0 | - | - |
| 45 | 53 | M | C | 0 | - | - |
| 46 | 78 | F | C | 0 | - | - |
| 47 | 76 | M | N | 0 | - | - |
| 48 | 71 | M | C | 1 | - | - |

| | | | | | | |
|---|---|---|---|---|---|---|
| 49 | 66 | F | C | 0 | - | - |
| 50 | 64 | F | C | 0 | - | - |
| 51 | 64 | M | C | 0 | - | - |
| 52 | 74 | M | N | 0 | - | - |
| 53 | 74 | F | C | 1 | - | - |
| 54 | 71 | F | C | 1 | - | - |
| 55 | 71 | F | C | 0 | - | - |
| 56 | 70 | F | C | 0 | - | - |
| 57 | 70 | F | C | 1 | - | - |
| 58 | 65 | M | C | 1 | - | - |
| 59 | 64 | F | F | 0 | - | - |
| 60 | 63 | M | N | 1 | - | - |
| 61 | 62 | M | F | 0 | - | - |
| 62 | 61 | F | C | 1 | - | - |
| 63 | 58 | M | C | 0 | - | - |
| 64 | 58 | M | C | 1 | - | - |
| 65 | 57 | M | C | 0 | - | - |
| 66 | 56 | F | C | 0 | - | - |
| 67 | 56 | M | C | 1 | - | - |
| 68 | 77 | M | C | 1 | - | - |
| 69 | 77 | M | F | 1 | - | - |
| 70 | 65 | F | F | 1 | - | - |
| 71 | 67 | M | C | 2 | - | - |
| 72 | 56 | F | N | 0 | - | - |
| 73 | 79 | F | F | 1 | - | - |
| 74 | 76 | F | C | 0 | - | - |
| 75 | 74 | M | F | 1 | - | - |
| 76 | 69 | F | F | 1 | - | - |
| 77 | 66 | M | F | 1 | - | - |
| 78 | 65 | M | F | 0 | - | - |
| 79 | 64 | M | F | 0 | - | - |
| 80 | 56 | M | F | 1 | - | - |
| 81 | 56 | M | C | 0 | - | - |
| 82 | 78 | M | F | 1 | - | - |
| 83 | 70 | F | N | 1 | - | - |
| 84 | 66 | F | N | 0 | - | - |
| 85 | 65 | M | C | 0 | - | - |
| 86 | 62 | M | F | 0 | - | - |
| 87 | 60 | F | N | 1 | - | - |
| 88 | 57 | M | F | 0 | - | - |
| 89 | 76 | F | C | 0 | - | - |
| 90 | 62 | M | C | 0 | - | - |
| 91 | 70 | F | F | 1 | - | - |
| 92 | 67 | M | F | 0 | - | - |
| 93 | 76 | F | F | 0 | - | - |
| 94 | 81 | F | N | 1 | - | - |
| 95 | 79 | F | N | 1 | - | - |
| 96 | 72 | M | C | 1 | - | - |
| 97 | 69 | M | C | 1 | - | - |
| 98 | 62 | F | C | 1 | - | - |
| 99 | 62 | M | C | 0 | - | - |
| 100 | 61 | F | C | 0 | - | - |
| 101 | 59 | F | F | 0 | - | - |
| 102 | 58 | M | F | 0 | - | - |
| 103 | 83 | F | N | 1 | - | - |
| 104 | 77 | F | C | 1 | - | - |
| 105 | 76 | F | F | 1 | - | - |
| 106 | 58 | M | C | 1 | - | - |
| 107 | 76 | M | N | 1 | - | - |
| 108 | 75 | M | C | 0 | - | - |
| 109 | 74 | M | F | 0 | - | - |

| | | | | | | |
|---|---|---|---|---|---|---|
| 110 | 70 | F | F | 2 | - | - |
| 111 | 67 | M | F | 0 | - | - |
| 112 | 63 | M | C | 1 | - | - |
| 113 | 71 | M | C | 1 | - | - |
| 114 | 51 | M | C | 0 | - | - |
| 115 | 61 | M | C | 1 | - | - |
| 116 | 78 | M | C | 0 | - | - |
| 117 | 60 | F | C | 0 | - | - |
| 118 | 74 | F | F | 0 | - | - |
| 119 | 71 | F | N | 0 | - | - |
| 120 | 80 | M | F | 0 | - | - |
| 121 | 62 | M | F | 1 | - | - |
| 122 | 75 | M | C | 2 | - | - |
| 123 | 69 | F | C | 1 | - | - |
| 124 | 70 | F | F | 0 | - | - |
| 125 | 67 | M | C | 1 | - | - |
| 126 | 74 | F | C | 0 | - | - |
| 127 | 55 | M | F | 0 | - | - |
| 128 | 59 | M | F | 0 | - | - |
| 129 | 73 | M | F | 0 | - | - |
| 130 | 64 | F | C | 0 | - | - |
| 131 | 63 | M | C | 0 | - | - |
| 132 | 80 | F | F | 0 | - | - |
| 133 | 72 | M | C | 1 | - | - |
| 134 | 75 | M | F | 1 | - | - |
| 135 | 54 | M | C | 0 | - | - |
| 136 | 68 | M | F | 0 | - | - |
| 137 | 55 | F | C | 0 | - | - |
| 138 | 68 | F | F | 1 | - | - |
| 139 | 70 | F | C | 1 | - | - |
| 140 | 79 | F | F | 1 | - | - |
| 141 | 61 | F | C | 0 | - | - |
| 142 | 74 | F | C | 1 | - | - |
| 143 | 76 | F | C | 0 | - | - |
| 144 | 63 | M | N | 0 | - | - |
| 145 | 60 | F | C | 0 | - | - |
| 146 | 65 | F | C | 1 | - | - |
| 147 | 69 | F | C | 0 | - | - |
| 148 | 70 | M | C | 1 | - | - |
| 149 | 77 | F | C | 0 | - | - |
| 150 | 73 | F | N | 1 | - | - |
| 151 | 61 | M | F | 0 | - | - |
| 152 | 65 | M | F | 1 | - | - |
| 153 | 69 | M | N | 0 | - | - |
| 154 | 67 | M | C | 1 | - | - |
| 155 | 66 | F | F | 1 | - | - |
| 156 | 67 | M | F | 0 | - | - |
| 157 | 67 | M | C | 1 | - | - |
| 158 | 80 | F | C | 1 | - | - |
| 159 | 63 | F | C | 1 | - | - |
| 160 | 69 | F | N | 0 | - | - |
| 161 | 62 | F | N | 1 | - | - |
| 162 | 82 | M | F | 1 | - | - |
| 163 | 75 | F | F | 1 | - | - |
| 164 | 62 | M | F | 0 | - | - |
| 165 | 58 | M | C | 0 | - | - |
| 166 | 60 | F | C | 0 | - | - |
| 167 | 65 | F | F | 1 | - | - |
| 168 | 66 | M | C | 0 | - | - |
| 169 | 66 | M | F | 0 | - | - |
| 170 | 62 | F | C | 0 | - | - |

| | | | | | | |
|---|---|---|---|---|---|---|
| 171 | 68 | M | C | 0 | - | - |
| 172 | 61 | F | C | 0 | - | - |
| 173 | 69 | F | F | 0 | - | - |
| 174 | 76 | F | C | 1 | - | - |
| 175 | 70 | F | N | 0 | - | - |
| 176 | 74 | F | F | 1 | - | - |
| 177 | 76 | F | C | 0 | - | - |
| 178 | 70 | M | C | 0 | - | - |
| 179 | 65 | F | F | 1 | - | - |
| 180 | 51 | F | N | 0 | - | - |
| 181 | 62 | F | C | 0 | - | - |
| 182 | 60 | M | C | 0 | - | - |
| 183 | 64 | F | F | 0 | - | - |
| 184 | 77 | F | C | 0 | - | - |
| 185 | 71 | F | F | 0 | - | - |
| 186 | 61 | F | F | 0 | - | - |
| 187 | 57 | M | C | 0 | - | - |
| 188 | 69 | M | N | 1 | - | - |
| 189 | 75 | M | C | 1 | - | - |
| 190 | 74 | F | C | 0 | - | - |
| 191 | 65 | F | C | 0 | - | - |
| 192 | 60 | F | C | 0 | - | - |
| 193 | 68 | F | N | 0 | - | - |
| 194 | 60 | M | N | 0 | - | - |
| 195 | 52 | M | C | 0 | - | - |
| 196 | 73 | F | C | 1 | - | - |
| 197 | 68 | F | F | 1 | - | - |
| 198 | 65 | M | C | 1 | - | - |
| 199 | 71 | M | C | 0 | - | - |
| 200 | 75 | M | F | 0 | - | - |
| 201 | 69 | M | C | 1 | - | - |
| 202 | 69 | M | C | 0 | - | - |
| 203 | 72 | F | F | 0 | - | - |
| 204 | 68 | F | C | 1 | - | - |
| 205 | 61 | M | C | 0 | - | - |
| 206 | 56 | M | C | 0 | - | - |
| 207 | 69 | F | C | 1 | - | - |
| 208 | 66 | M | F | 0 | - | - |
| 209 | 77 | F | C | 1 | - | - |
| 210 | 63 | M | C | 1 | - | - |
| 211 | 60 | M | F | 0 | **+** | **+** |
| 212 | 67 | M | C | 0 | - | - |
| 213 | 67 | F | C | 1 | - | - |
| 214 | 64 | F | C | 0 | - | - |
| 215 | 76 | M | F | 0 | - | - |
| 216 | 77 | M | F | 1 | - | - |
| 217 | 56 | F | C | 0 | - | - |
| 218 | 74 | M | F | 1 | - | - |
| 219 | 59 | F | F | 0 | - | - |
| 220 | 73 | M | F | 1 | - | - |
| 221 | 68 | M | C | 0 | - | - |
| 222 | 75 | F | C | 0 | - | - |
| 223 | 67 | M | C | 0 | - | - |
| 224 | 69 | M | F | 1 | - | - |
| 225 | 74 | F | F | 1 | - | - |
| 226 | 49 | F | C | 0 | - | - |
| 227 | 64 | M | C | 0 | - | - |
| 228 | 64 | F | N | 0 | - | - |
| 229 | 75 | M | F | 1 | - | - |
| 230 | 69 | F | N | 0 | - | - |
| 231 | 64 | F | C | 0 | - | - |

| | | | | | | |
|---|---|---|---|---|---|---|
| 232 | 74 | F | F | 0 | - | - |
| 233 | 60 | M | C | 0 | - | - |
| 234 | 49 | F | C | 1 | - | - |
| 235 | 62 | M | C | 0 | - | - |
| 236 | 59 | F | C | 0 | - | - |
| 237 | 65 | M | F | 0 | - | - |
| 238 | 61 | M | F | 0 | - | - |
| 239 | 82 | M | C | 1 | - | - |
| 240 | 72 | M | F | 0 | - | - |
| 241 | 60 | M | F | 0 | - | - |
| 242 | 84 | M | F | 1 | - | - |
| 243 | 65 | F | C | 1 | - | + |
| 244 | 80 | M | F | 1 | - | - |
| 245 | 69 | F | F | 0 | - | - |
| 246 | 58 | M | F | 1 | - | - |
| 247 | 66 | M | F | 1 | - | - |
| 248 | 59 | M | F | 1 | - | - |
| 249 | 67 | M | F | 1 | - | - |
| 250 | 60 | M | F | 0 | - | - |
| 251 | 76 | M | F | 0 | - | - |
| 252 | 68 | F | C | 0 | - | - |
| 253 | 75 | F | N | 0 | - | - |
| 254 | 64 | F | F | 1 | - | - |
| 255 | 62 | M | F | 1 | - | - |
| 256 | 72 | M | N | 0 | - | - |
| 257 | 77 | M | C | 0 | - | - |
| 258 | 65 | F | F | 0 | - | - |
| 259 | 75 | F | C | 0 | - | - |
| 260 | 67 | M | C | 1 | - | - |
| 261 | 66 | F | C | 1 | - | - |
| 262 | 73 | F | F | 0 | - | - |
| 263 | 55 | F | C | 1 | - | - |
| 264 | 55 | M | F | 0 | - | - |
| 265 | 78 | F | N | 1 | - | - |
| 266 | 67 | F | F | 1 | - | - |
| 267 | 54 | F | C | 1 | - | - |
| 268 | 77 | M | F | 1 | - | - |
| 269 | 74 | M | C | 0 | - | - |
| 270 | 66 | F | C | 1 | - | - |
| 271 | 65 | F | C | 0 | - | - |
| 272 | 67 | F | C | 0 | - | - |
| 273 | 66 | M | C | 0 | - | - |
| 274 | 59 | M | F | 0 | - | - |
| 275 | 63 | F | C | 1 | - | - |
| 276 | 76 | M | F | 1 | - | - |
| 277 | 60 | M | C | 0 | - | - |
| 278 | 66 | F | C | 0 | - | - |
| 279 | 62 | F | C | 0 | - | - |
| 280 | 62 | M | C | 1 | - | - |
| 281 | 68 | F | C | 1 | - | - |
| 282 | 77 | M | C | 1 | - | - |
| 283 | 63 | M | C | 0 | - | - |
| 284 | 64 | M | C | 1 | - | - |
| 285 | 48 | F | F | 1 | - | - |
| 286 | 72 | M | C | 0 | - | - |
| 287 | 64 | F | C | 1 | - | - |
| 288 | 67 | F | C | 0 | - | - |
| 289 | 57 | F | N | 0 | - | - |
| 290 | 73 | M | F | 0 | - | - |
| 291 | 79 | M | F | 1 | - | - |
| 292 | 73 | F | F | 0 | - | - |

| | | | | | | |
|---|---|---|---|---|---|---|
| 293 | 76 | F | F | 0 | - | - |
| 294 | 67 | M | F | 1 | - | - |
| 295 | 67 | F | C | 1 | - | - |
| 296 | 73 | F | F | 0 | - | - |
| 297 | 60 | F | F | 0 | - | - |
| 298 | 64 | F | N | 1 | - | - |
| 299 | 65 | F | C | 0 | - | - |
| 300 | 70 | M | C | 1 | - | - |
| 301 | 79 | F | F | 1 | - | - |
| 302 | 45 | F | N | 0 | - | - |
| 303 | 70 | M | F | 0 | - | - |
| 304 | 68 | F | N | 0 | - | - |
| 305 | 76 | M | N | 0 | - | - |
| 306 | 58 | M | F | 0 | - | - |
| 307 | 62 | F | C | 0 | - | - |
| 308 | 65 | F | F | 0 | - | - |
| 309 | 51 | F | C | 0 | - | - |
| 310 | 69 | F | F | 0 | - | - |
| 311 | 57 | F | F | 0 | - | - |
| 312 | 71 | F | C | 1 | - | - |
| 313 | 74 | M | F | 1 | - | - |
| 314 | 62 | M | C | 1 | - | - |
| 315 | 76 | F | F | 0 | - | - |
| 316 | 62 | F | F | 0 | - | - |
| 317 | 61 | M | F | 0 | - | - |
| 318 | 54 | F | N | 0 | - | - |
| 319 | 42 | F | N | 0 | - | - |
| 320 | 66 | M | F | 1 | - | - |
| 321 | 60 | F | C | 0 | - | - |
| 322 | 74 | F | F | 0 | - | - |
| 323 | 77 | M | C | 1 | - | - |
| 324 | 70 | M | F | 1 | - | - |
| 325 | 64 | M | C | 0 | - | - |
| 326 | 67 | F | C | 0 | - | - |
| 327 | 62 | F | C | 0 | - | - |
| 328 | 74 | M | F | 1 | - | - |
| 329 | 69 | M | F | 1 | - | - |
| 330 | 69 | M | F | 1 | - | - |
| 331 | 69 | F | C | 0 | - | - |
| 332 | 74 | M | F | 0 | - | - |
| 333 | 77 | M | C | 0 | - | - |
| 334 | 67 | F | F | 0 | - | - |
| 335 | 67 | F | N | 0 | - | - |
| 336 | 67 | M | N | 1 | - | - |
| 337 | 66 | F | F | 1 | - | - |
| 338 | 75 | F | C | 0 | - | - |
| 339 | 66 | F | C | 0 | - | - |
| 340 | 62 | F | C | 0 | - | - |
| 341 | 61 | F | C | 0 | - | - |
| 342 | 74 | M | F | 0 | - | - |
| 343 | 71 | F | F | 1 | - | - |
| 344 | 62 | F | C | 0 | - | - |
| 345 | 56 | F | C | 0 | - | - |
| 346 | 69 | F | C | 0 | - | - |
| 347 | 72 | F | C | 0 | - | - |
| 348 | 70 | F | C | 0 | - | - |
| 349 | 69 | M | C | 1 | - | - |
| 350 | 75 | M | C | 1 | - | - |
| 351 | 75 | F | C | 1 | - | - |
| 352 | 70 | M | C | 0 | - | - |
| 353 | 71 | F | F | 0 | - | - |

| | | | | | | |
|---|---|---|---|---|---|---|
| 354 | 73 | M | C | 0 | - | - |
| 355 | 71 | M | C | 0 | - | - |
| 356 | 62 | F | C | 0 | - | - |
| 357 | 71 | M | F | 0 | - | - |
| 358 | 77 | M | F | 0 | - | - |
| 359 | 64 | F | C | 0 | - | - |
| 360 | 56 | F | C | 1 | - | - |
| | Mean (67.1 years) | Female (50.8%), Male (49.2%) | C (50.8%), F (37.5%), N (11.7%) | PS 0 (60.0%), PS 1 (38.6%), PS 2 (1.4%) | Positive cases (0.3%) | Positive cases (0.6%) |
